# The subjective amplitude of the diurnal rhythm matters - chronobiological insights for neuroimaging studies

**DOI:** 10.1101/2023.05.17.541153

**Authors:** Michal Rafal Zareba, Patrycja Scislewska, Magdalena Fafrowicz, Tadeusz Marek, Halszka Oginska, Iwona Szatkowska, Ewa Beldzik, Aleksandra Domagalik

## Abstract

Multiple aspects of human physiology, including mood and cognition, are subjected to diurnal rhythms. While the previous neuroimaging studies have focused solely on the morningness-eveningness (ME) preference dichotomy, i.e. the circadian phase, the second key dimension of the diurnal rhythms, i.e. the strength of these preferences (amplitude; AM), has been completely overlooked. Uncovering the neural correlates of AM is especially important considering its link with negative emotionality. Structural T1-weighted neuroimaging data from 79 early (EC) and 74 late (LC) chronotypes were analysed to compare grey matter (GM) volume and cortical thickness. The study aimed to elucidate whether the subjective AM and its interaction with ME was a significant predictor of individual brain structure. Both GM volume and cortical thickness of the left primary visual cortex was negatively correlated with AM scores across the entire sample. Furthermore, EC and LC differed in their association between AM scores and the GM volume in the right middle temporal gyrus, with the positive and negative correlations reported respectively in the two groups. The current study underlines the importance of the visual system in circadian rhythmicity and provides possible neural correlates for AM-related differences in negative affect processing. Furthermore, the presence of the opposite correlations between brain anatomy and AM in the two groups suggests that the behavioural and neuronal chronotype differences might become more pronounced in individuals with extreme diurnal differences in mood and cognition, highlighting the necessity to additionally account for AM in neuroimaging studies.

**Highlights:**

- Structure of primary visual cortex is linked to subjective diurnal rhythms amplitude
- Middle temporal gyrus is sensitive to interaction of rhythm phase and distinctness
- Distinctness of the diurnal rhythms may modulate results of the neuroimaging studies

## 1. Introduction

According to the generally accepted concept of the endogenous biological clocks, the rhythmicity of the human body is based on the existence of multiple oscillators that receive afferent information about the environment (thus synchronising the body to the external conditions, e.g. the changes in time zones during travel) and subsequently transmit it to the body’s cells and tissues (thus influencing the rhythmicity of various processes) (Allada & Chung, 2010). One of the central elements of this brain machinery is the suprachiasmatic nucleus (SCN), primarily entrained with the visual information from the retina, a fact that underlines the importance of the light signals in the generation of the rhythms (Beytebiere et al., 2019; Duda et al., 2020; Reghunandanan & Reghunandanan, 2006).

In 1986, David Minors and James Waterhouse, researchers of human circadian rhythms, recognized that full description of the daily rhythms requires a characterisation of three distinct elements: phase (chronotype or morningness-eveningness, ME), stability and amplitude (Fisher, 1986). The ME score represents an individual’s preferred time of activity, while the amplitude score - the strength of this preference. On average, the endogenous circadian rhythm in humans has a period of about 24.2 hours with little variation from day to day (Czeisler & Gooley, 2007). However, the timing of the periods of rest and activity, i.e. ME, can vary greatly from person to person, making the individual chronotypes fall on a continuum, at the both ends of which there are individuals commonly referred to as “larks” and “owls”. The “larks”, or early chronotypes (EC), prefer to get up early in the morning and feel best in the early hours of the day. It is believed that their circadian periods may be shorter than 24 hours (Sack et al., 2007a). In contrast, the rhythms of “owls” or late chronotypes (LC) are characterised by periods that can be longer than 24 hours, which results in the peak of cognitive performance in the evening (Sack et al., 2007b). People who fall between these ends of the continuum are commonly referred to as intermediate types.

Population studies made it possible to determine that the probability distribution of chronotypes resembles the normal distribution with strong developmental changes in the daily preferences throughout life. The chronotype itself does not change during the life of individuals, but a shift in the preferred hours of activity is observed. In childhood, there is a tendency to show earlier time preferences. Adolescents go to bed late and tend to get up late the next day. However, as individuals get older, a gradual shift back towards the earlier times is observed. It is found that the subjective rhythm amplitude decreases with age, while stability of sleeping habits remains constant. (Di Milia & Folkard, 2021; Randler, 2016).

So far, the most common tools for the determination of one’s chronotype are the Morningness-Eveningness Questionnaire (MEQ) (Horne & Ostberg, 1976) and Munich Chronotype Questionnaire (MCTQ) (Roenneberg et al., 2015). They allow measuring the “phase” (morningness-eveningness, ME) of diurnal rhythms and are used in most circadian studies. However, these tools overlook the subjective amplitude (AM), i.e. the distinctness of the rhythm. Unlike the traditional classification of chronotypes, which determines the type of “extreme” or “moderate” phenotypes based on the exact hours of waking up and falling asleep, the AM refers to the perceived intensity of differences in functioning between the morning and evening hours. Thus, a “definitely morning” person is one who naturally gets up early and quickly reaches maximum arousal, but in the evening, when it is their usual bedtime, they tend not to be able to control the drowsiness. In turn, a “moderately morning” person is one who is in a better mood in the morning hours, but is also able to concentrate at any time of the day. (Kontrymowicz-Ogińska, 2011). These differences reflect individual sensitivity to the environmental changes; in terms of circadian rhythmicity, this would be to function ‘according to nature’, despite the masking influence of social circumstances. The characterisation of diurnal functioning in two dimensions - not only subjective phase (ME) but also amplitude (AM) - is consistent with the theory of biological rhythms. We believe that subjective AM reflects greater diurnal variation in physiological processes, which in turn may be shaped by structural and functional specificities of the nervous system. Hence, the idea of looking for differences in the neural structures in individuals differing in sensitivity to synchronisers, i.e. showing different subjective AM of the circadian rhythm. Making such a distinction is especially important taking into account that a larger AM of diurnal variations in mood and cognition may be independently and more strongly related to negative emotionality than the eveningness preference (Carciofo, 2020, 2022; Nowakowska-Domagała et al., 2022). Current theories posit the social jetlag, i.e., a significant mismatch between the biologically preferred activity hours and the societally-demanded ones, as one of the main mechanisms linking circadian rhythms to mental health problems (Souêtre et al., 1989; Wittmann et al., 2006). People with more pronounced diurnal rhythms have more difficulties in adapting to these external requirements. Therefore, investigations probing the neural correlates of distinctness of the rhythms can greatly benefit our understanding of individual proneness for psychiatric disorders, shedding light on an important, although currently neglected factor. For this reason, the present work aimed to investigate how the AM of the diurnal variations in mood and cognition is related to structure of the brain, and whether these associations differ depending on individual chronotype.

Several previously published studies investigated the relationship between brain anatomy and ME in young adults. Amongst the most consistently reported brain structures, there are superior and inferior parietal lobules, precuneus and occipital lobes, which are characterised by higher grey matter (GM) volume and cortical thickness with increasing preference for eveningness (Evans et al., 2021; Rosenberg et al., 2018; Takeuchi et al., 2015). Additionally, greater cortical thickness in LC was reported in the inferior frontal gyrus pars triangularis, insula, as well as in the fusiform and entorhinal gyri (Rosenberg et al., 2018; Zareba et al., 2022b). In turn, an increased morningness preference has been linked to greater GM volume in the orbitofrontal cortex (Takeuchi et al., 2015). However, all the reports listed above omit the issue of the AM of diurnal rhythms, which may play a key role in the context of analysing individual differences. Thus, in the current work we set out to investigate how the GM volume and the cortical thickness are associated with the distinctness dimension of the circadian rhythms, and whether this relationship is modulated by individual chronotype. To the best of our knowledge, only one work has investigated whether the AM of circadian rhythms is related to the neuroimaging results (Farahani et al., 2022). It was reported that the graph theory measures derived on the basis of resting-state functional connectivity were indeed dependent on the rhythm distinctness across multiple brain networks. For this reason, we hypothesised that these findings could be at least in part reflected in the brain anatomy. Nevertheless, the aforementioned study did not test for the interactions between the circadian phase and the perceived rhythm amplitude. Consequently, an exploratory analysis was performed to fill this crucial gap in the literature.

## 2. Methods

### 2.1. Participants

The currently described analyses were performed using a fully anonymised high resolution structural dataset obtained during fMRI projects (N = 153). The majority of this data has been made freely available to other scientists (Zareba et al., 2022a). Compared to the published dataset, the present investigation covers additional 17 participants, which were not included in the aforementioned publication due to the lack of full sleep-related data. All subjects were aged from 19 to 35, right-handed, had normal or corrected-to-normal vision, were drug-free, and self-reported no history of either neurological, nor psychiatric disorders. Trait-like chronotype was assessed using of the Chronotype Questionnaire (ChQ) (Ogińska, 2011; Oginska et al., 2017). The tool is particularly useful for distinguishing between the two dimensions of circadian rhythms, i.e. the subjective circadian phase (ME) and the subjective amplitude of diurnal variations in mood and cognition (AM). Each dimension is tested with 8 objects ranked from 1 to 4, which are summed together, making the possible score range from 8 to 32 points. Example questions probing ME are the following: “I feel sluggish in the morning and I warm up slowly during the day” and “I am usually in an excellent mood in the morning”. In turn, AM is tested with items like “I feel substantial variations of my mood during the day” and “When my usual sleep time comes, I can hardly overcome sleepiness”. The higher the score in the ME subscale, the more evening-oriented the individual is. Similarly, the higher the score in the AM subscale, the stronger the within-day fluctuations in mental processes.

In the current dataset, the ME scores ranged from 11 to 32 and were not distributed normally according to the Kolmogorov-Smirnov test (p < 0.05). As such, the sample was divided into two groups: early chronotypes (EC; ME scores from 11 to 22) and late chronotypes (LC; ME scores from 23 to 32). The scores in AM varied from 11 to 29 and followed the normal distribution according to the Kolmogorov-Smirnov test (p > 0.05). The EC and LC groups did not differ in the AM scores (p = 0.13). The demographic summary of the sample is presented in Table 1.

**Table 1.** Demographic summary of the cohort used in the current project.

| Variable | EC, n = 79<br>(mean ± SD) | LC, n = 74<br>(mean ± SD) | Group<br>difference<br>(p value) | Full cohort, n = 153<br>(mean ± SD) |
| --- | --- | --- | --- | --- |
| Sex (male/female) | 28/51 | 26/48 | NA | 54/99 |
| Age (years) | 25.06 ± 4.17 | 23.72 ± 3.40 | 0.030 <sup>NP</sup> | 24.42 ± 3.87 |
| ME | 17.03 ± 4.03 | 27.16 ± 2.56 | < 0.001 <sup>NP</sup> | 21.93 ± 5.77 |
| AM | 20.61 ± 4.03 | 21.51 ± 3.33 | 0.133 <sup>P</sup> | 21.05 ± 3.72 |
| PSQI | 2.92 ± 1.29 <sup>M</sup> | 3.22 ± 1.53 <sup>M</sup> | 0.389 <sup>NP</sup> | 3.06 ± 1.41 <sup>M</sup> |
| ESS | 6.86 ± 3.17 <sup>M</sup> | 7.34 ± 3.88 <sup>M</sup> | 0.443 <sup>P</sup> | 7.09 ± 3.52 <sup>M</sup> |

Abbreviations: EC = early chronotype; LC = late chronotype; SD = standard deviation; NA = non-applicable; ME = morningness-eveningness scale from the Chronotype Questionnaire (ChQ) (Ogińska, 2011; Oginska et al., 2017); AM = amplitude scale from ChQ; PSQI = Pittsburgh Sleep Quality Index (Buysse et al., 1989); ESS = Epworth Sleepiness Scale (Johns, 1991).
<sup>NP</sup> Assessed with a non-parametric Mann-Whitney U test.
<sup>P</sup> Assessed with a parametric two sample t-test.
<sup>M</sup> Data not available for 7 EC and 9 LC.

### 2.2. MRI data acquisition

Magnetic resonance imaging (MRI) was performed on a 3T scanner (Magnetom Skyra, Siemens) using a 20-channel or 64-channel head/neck coil. The data was collected with the MPRAGE sequence (176 sagittal slices; 1×1×1.1 mm voxel size; TR = 2300 ms, TE = 2.98 ms, flip angle = 9°, GRAPPA acceleration factor 2). All scanning sessions were performed between 4:30 PM and 8:55 PM to minimise the effects of time-of-day on the morphometric measures (Trefler et al. 2016).

### 2.3 Voxel-based morphometry

Voxel-based morphometry (VBM) analysis was performed in the CAT12 toolbox (Gaser & Kurth, 2021). The brain images were segmented into GM, white matter, and cerebrospinal fluid, and non-linearly normalised in the MNI152 space using diffeomorphic anatomical registration through exponentiated lie algebra (DARTEL). The normalised GM volumes were modulated with the Jacobian determinant to account for the resulting volumetric changes. At this point, the GM segments of all the subjects were inspected visually to ensure the satisfactory quality of the preprocessing. This served as a further confirmation of the routine quality reports created by the software. As a result of this double check procedure, data from all the participants was included in the further steps of the analyses. The normalised and modulated GM segments were smoothed with a 4 mm Gaussian filter. The statistical analysis was performed in the 3dMVM program (Chen et al., 2014) in AFNI (Cox, 1996). The model aimed to investigate whether AM and its interaction with ME was a significant predictor of the variability in GM volume. Sex, age and total intracranial volume of the participants were controlled as covariates. The multiple comparisons correction was achieved with the cluster-level family-wise error correction (FWE < 0.05) following the initial voxel-level thresholding (p < 0.001).

### 2.4. Cortical thickness

Estimation of cortical thickness was achieved using the projection-based method in the CAT12 toolbox (Gaser & Kurth, 2021). After the brain tissue segmentation, the white matte distance was calculated and the local maxima were projected onto GM voxels with the use of a neighbouring relationship described by the white matter distance. Thanks to this procedure, the preprocessing pipeline was capable of handling sulcal asymmetries, sulcal blurring and partial voluming effects without the explicit reconstruction of sulcus. The next steps of the stream included topology correction, spherical mapping and spherical registration (Yotter et al., 2011). The normalised cortical thickness maps were smoothed with a 8 mm Gaussian filter. The statistical analysis was performed in CAT12 (Gaser & Kurth, 2021). Similarly to the VBM analysis, the contrasts of interest examined whether cortical thickness was associated with AM, as well as its interaction with ME. Sex and age were included in the model as covariates. The multiple comparison correction was achieved through the cluster-level FWE (FWE < 0.05) after the vertex-wise thresholding (p < 0.001).

## 3. Results

### 3.1. Voxel-based morphometry

VBM analysis revealed two significant clusters: one for the main effect of AM and one for the interaction between AM and ME (voxel-level p < 0.001, cluster-level FWE < 0.05). The GM volume in the left primary visual cortex (Brodmann Area (BA) 17) was negatively correlated with the individual AM scores (r = -0.396), independently of the ME grouping. In turn, the GM volume of the right middle temporal gyrus (BA21, BA37) had a differential association with AM scores depending on whether the individuals were classified as EC or LC. In the former group, the structural measure was positively linked to AM (r = 0.251) whereas in the latter group a negative relationship was found (r = -0.402). The results are presented in more detail in Table 2, as well as Figures 1 and 2.

**Table 2.**
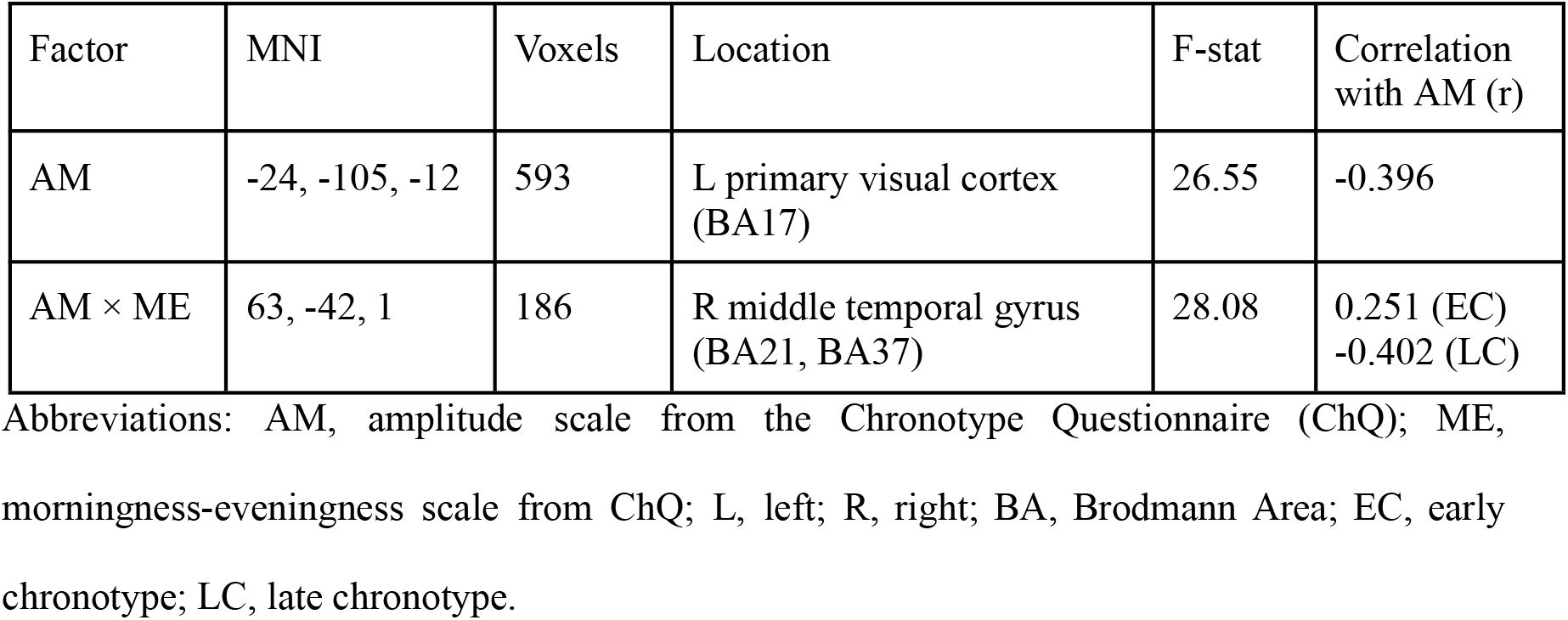
Results of the voxel-based morphometry analysis (voxel-level p < 0.001, cluster-level FWE < 0.05).

**Figure 1.**
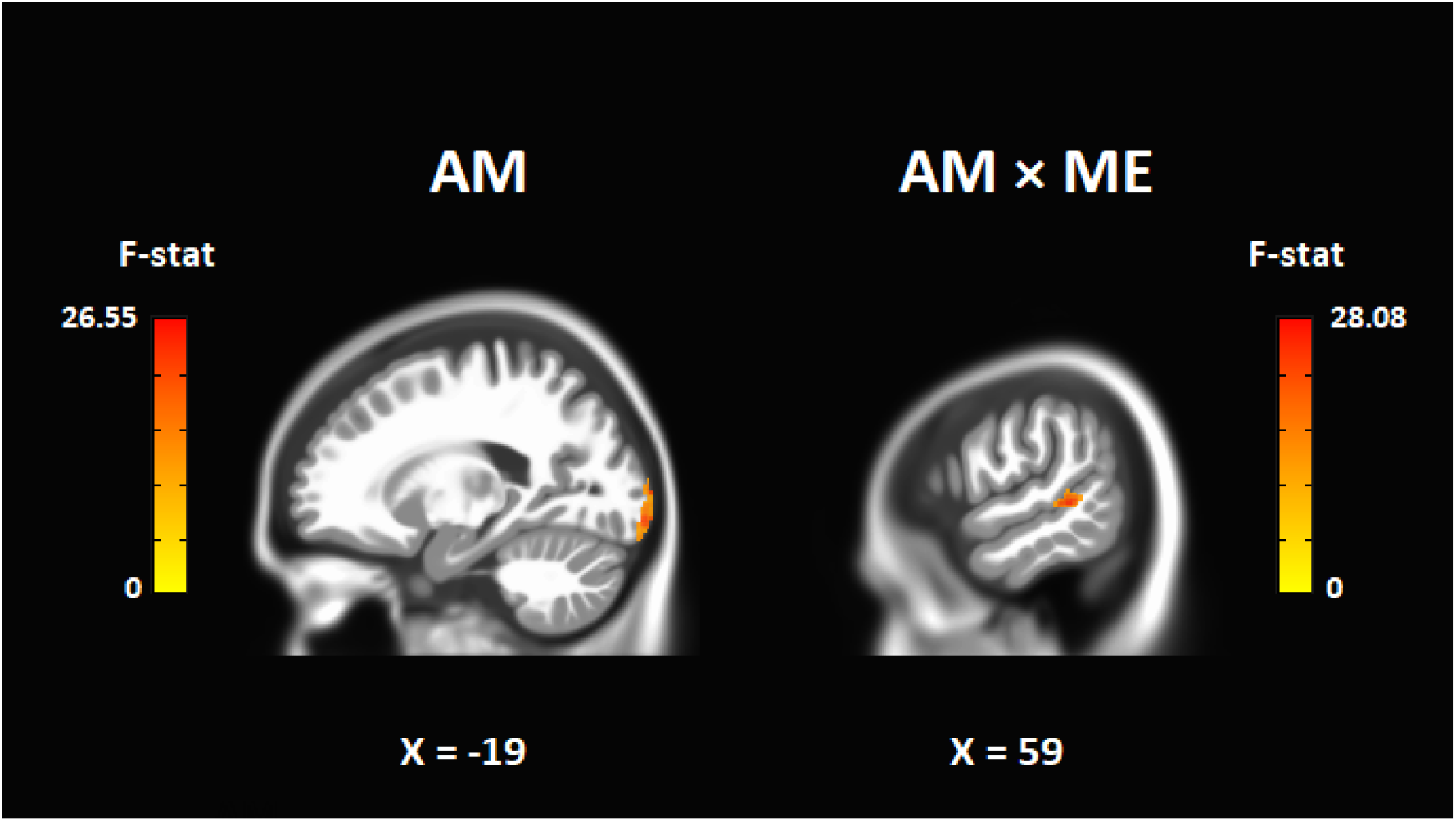
Grey matter volume in the left primary visual cortex was negatively associated with the subjective amplitude of diurnal variations (AM) regardless of chronotype (ME). An interaction between AM and ME was found in the right middle temporal gyrus.

**Figure 2.**
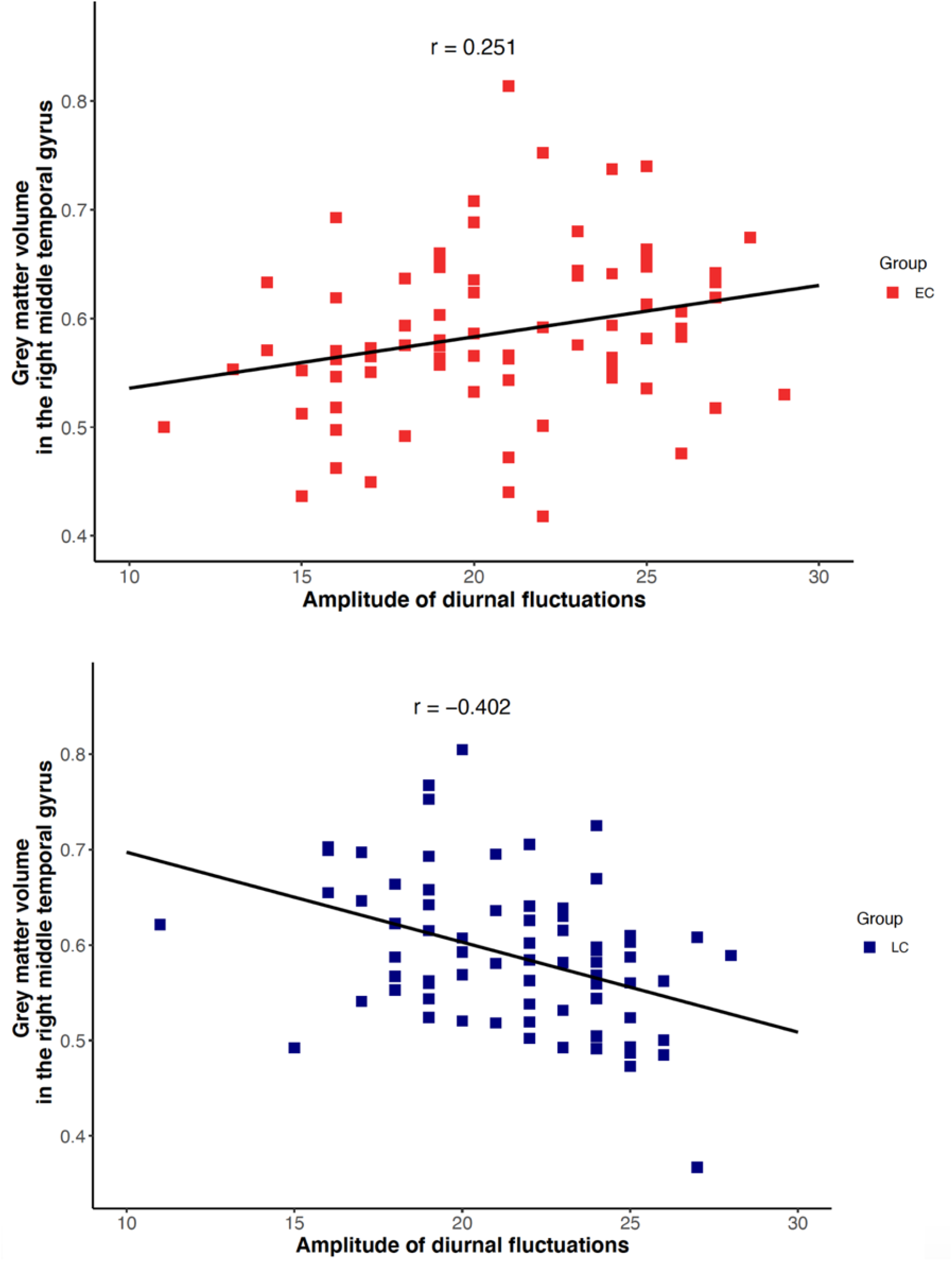
Early (EC) and late chronotypes (LC) differed in their relationship between the grey matter volume in the right middle temporal gyrus and the subjective amplitude of diurnal fluctuations. While in the case of EC the relationship was positive (top panel; r = 0.251), a negative association was found for LC (bottom panel; r = -0.402).

### 3.2. Cortical thickness

The analysis performed at the threshold of vertex-level p < 0.001 and cluster-level FEW < 0.05 revealed no significant findings for both the direct effect of AM and its interaction with ME. However, an exploratory analysis with a slightly less stringent thresholding (vertex-level p < 0.005 and cluster-level FWE < 0.05) revealed that across the entire sample AM scores were negatively correlated with the cortical thickness of the left primary visual cortex (BA17), mirroring the VBM findings. The results are described in more detail in Table 3 and Figure 3.

**Table 3.** Results of the cortical thickness analysis (vertex-level p < 0.005, cluster-level FWE < 0.05).

| Factor | Coordinates | Vertices | Location | T-stat | Correlation with AM (r) |
| --- | --- | --- | --- | --- | --- |
| AM | -17, -101, -9 | 102 | L primary visual cortex (BA17) | -4.24 | -0.309 |

Abbreviations: AM, amplitude scale from the Chronotype Questionnaire (ChQ); L, left; BA, Brodmann Area.

**Figure 3.**
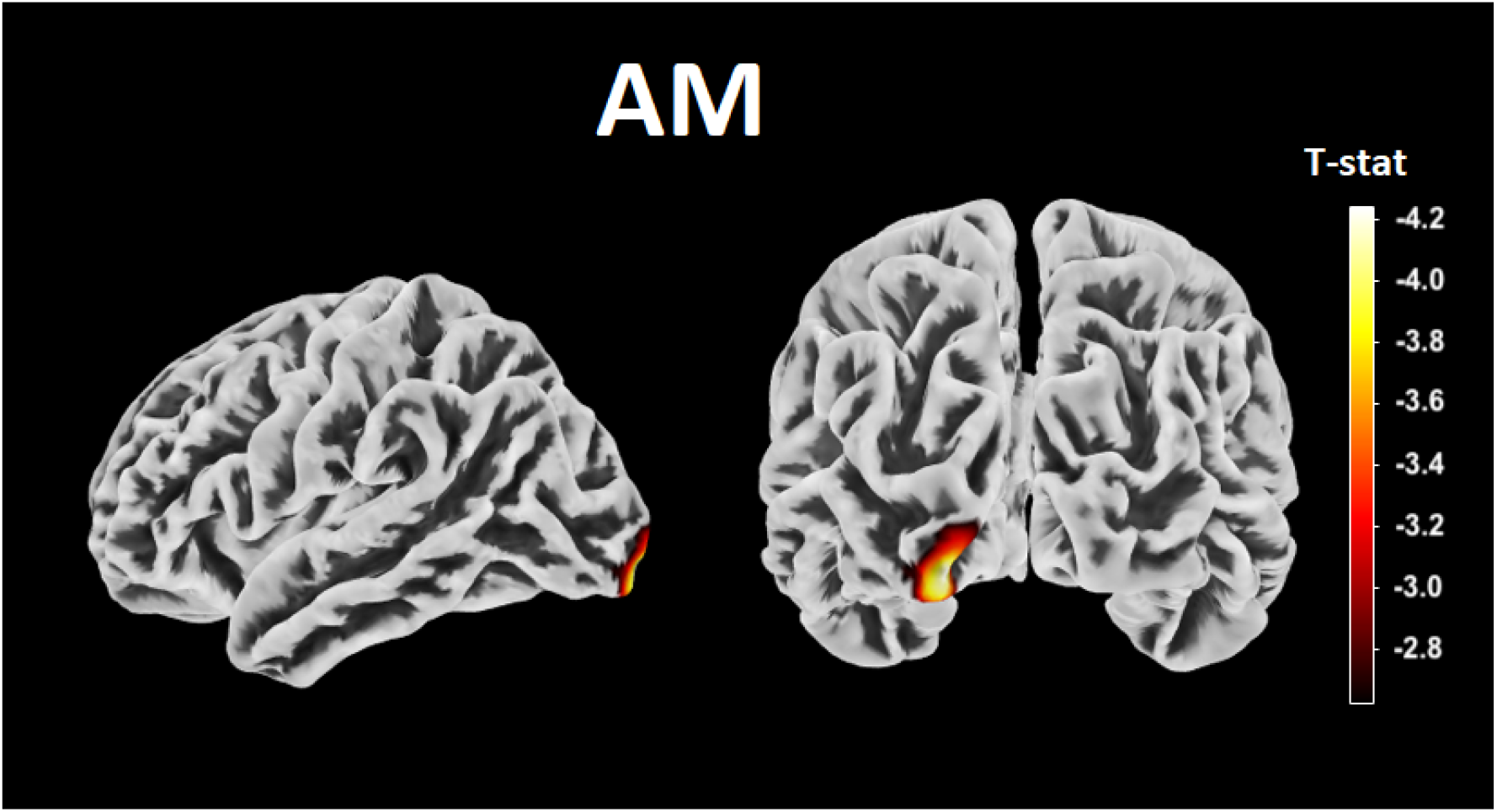
Subjective amplitude of diurnal variations (AM) in mood and cognition was negatively correlated with the cortical thickness of the left primary visual cortex (vertex-level p < 0.005, cluster-level FWE < 0.05).

## 4. Discussion

A few neuroimaging studies published so far have investigated how indices of brain structure and function are related to the ME aspect of the circadian rhythms in humans. Despite the fact that AM dimension is an equally important factor describing the functioning of the individual biological clock, this subject is yet to receive attention from a wider audience of scientists working at the border of cognitive neuroscience and chronobiology. Thus, the current work makes a significant advancement in the field by reporting for the first time that the macroscopic brain structure is related both to the AM of the rhythms, as well as its interactions with the ME dimension.

The study has found a negative correlation between the GM volume and cortical thickness of the left primary visual cortex and AM (respectively r = -0.396 and r = -0.309). This means that no matter what the subject’s chronotype is, the more pronounced the biological rhythm (and thus the greater the subjective amplitude), the less GM in the left primary visual cortex. Such a consistency of the results pinpoints the importance of the relationship between the structure of the early visual system and the strength of daily fluctuations in mood and cognition. As high AM of the rhythms is related to negative affectivity (Staller & Randler 2021), we believe that the relationship between high AM and smaller primary visual cortex in the left hemisphere may reflect deficient regulation of negative emotions. Our findings correspond anatomically with the established theories on how light can exert its impact on one’s brain. Light acts as one of the primary zeitgebers determining the human circadian rhythms and this information is largely relayed through the system associated with melanopsin-containing intrinsically-photosensitive retinal ganglion cells (ipRGCs) (Duda et al., 2020). Light stimulation targeting this specific class of photoreceptors has been reported to decrease the alpha band activity in occipital areas similar to those reported here (Vandewalle et al., 2018) Furthermore, the ipRGCs-related system exerts a pronounced modulatory effect on mood and cognition (An et al., 2020; Fernandez et al., 2018), however, the affective contributions are likely more mediated by its subcortical rather than cortical elements (Gaggioni et al., 2014). On the other hand, the primary visual cortex in the left hemisphere, together with the left orbitofrontal cortex, seems to belong to a neural network associated with regulation of negative emotions (Tak et al., 2021). Furthermore, the primary visual cortex seems to be involved in cognitive functions. For example, the individual differences in visual working memory (WM) storage may be predicted by the anatomical measures of the primary visual cortex volume, surface size, and cortical thickness, i.e., subjects with a bigger primary visual cortex have greater visual WM storage (Bergmann et al., 2016). Similarly, the sensory recruitment theory of working memory suggests a potential role for the early visual cortex in visual WM (Curtis and Sprague, 2021). One emerging view is that information in WM is maintained through sensory recruitment so that information is stored through continuous activity in sensory areas that encode the information to be remembered (Serences et al., 2009). What is more, Harrison and Tong (2009) proved that early visual areas can retain specific information about visual features stored in working memory for many seconds, even when there is no physical stimulus. Further functional neuroimaging research should clarify whether the relationships observed in our study may reflect the AM-related differences in the processing of negative emotions. Additionally, given the association between the structure of the primary visual cortex and visual WM efficiency (Bergmann et al., 2016), it would also be interesting to investigate whether the performance in this cognitive domain is related to the AM of diurnal rhythms.

Interestingly, the only published study to investigate the relationship between AM and neuroimaging indices reported that in the late afternoon and evening (i.e.the same scan timing as in our investigation), the resting-state functional connectivity-derived graph theory measures in the early visual cortices were related to individual AM scores, showing, in conjunction with our results, that this dimension of circadian rhythms is related to both the structure and function of the occipital areas (Farahani et al., 2022). When comparing our findings with the reports describing the anatomical brain correlates of the ME dimension of diurnal rhythms (Evans et al., 2021; Rosenberg et al., 2018; Takeuchi et al., 2015), AM is associated with the variability in the most posterior and ventral parts of the occipital lobes, while differences in ME are reflected in more anterior and dorsal areas of this region. When combined, these findings underline the important link between the visual system and circadian rhythmicity.

Our investigation further reports the dependence of the GM volume in the right middle temporal gyrus on the interaction between AM and ME. In the case of EC, there is a weak positive correlation with AM (r = 0.251), while in the case of LC – moderate negative (r = -0.402). According to the Human Brainnetome Atlas (http://atlas.brainnetome.org), the highlighted region is implicated in both the social cognition-related processes, such as action observation or face and tone monitoring, and reward-related functions, namely delay discounting (Fan et al., 2016). Both these domains are known to be affected by the ME dimension of diurnal rhythms. LC individuals are characterised by increased negative or decreased positive processing in tasks probing categorisation, recognition, and recall of emotions, independently of sleep problems (Berdynaj et al., 2016; C. M. Horne et al., 2017). Furthermore, they show an increased drive towards smaller immediate rewards over delayed larger ones and decreased brain activity in the medial prefrontal cortex during reward anticipation (S. L. Evans & Norbury, 2021; Hasler et al., 2013; Stolarski et al., 2013). The fact that the volume of the right middle temporal gyrus has differential association with AM in the two groups suggests that these behavioural and neuronal differences might become more pronounced in individuals with both extremely low and high values of perceived diurnal differences in mood and cognition. This, in turn, raises an important question in relation to the previous studies. If the AM dimension is not accounted for as one of the factors, it may lead to an unwanted bias in the reported results. An extreme owl with a very distinct rhythm may be characterised by different features than an owl with a less pronounced biological rhythm. For this reason, we underline the importance of considering AM as a factor potentially modulating the results of all studies on chronotypes.

An important and quite difficult issue to consider in circadian rhythms research is the very close connection between chronotypes and sleep quality. Especially in the case of LC, it is difficult to state to what extent the differences in behaviour or neuroanatomy are affected by shortened sleep duration or problematic wake-up times. To exclude the possibility that the current sleeping problems affect the results, in our study EC and LC groups do not differ in PSQI and ESS scores (p > 0.05). Furthermore, the GM volume and the cortical thickness in the significant clusters is not correlated with the scores in any of these scales (p > 0.05), which confirms that these findings are indeed related to AM and its interaction with ME. One limitation of the current study, however, is that PSQI and ESS scores were not available for 16 out of 153 participants. Secondly, EC and LC groups slightly differed in their mean age (25.06 ± 4.17 vs. 23.72 ± 3.40), however, this effect was accounted for by including age as a covariate in the analyses. Last but not least, when considering neuroanatomical differences in people with different chronotypes, the cause-and-effect chain is often not clear. One can argue that the structural alterations of specific brain areas may predetermine one’s chronotype. Alternatively and equally likely, however, human behaviour resulting from the interaction of having a specific chronotype and the influence of environmental factors can affect the processes of neuroplasticity, which may, in turn, result in a change in the size of selected areas of the brain. Future longitudinal studies in young adults, similar to recent work in adolescents (Vulser et al., 2023), are thus warranted to elucidate the matter.

## 5. Conclusions

The current study extends the state of knowledge by showing that the subjective amplitude of circadian rhythms is an important factor that needs to be accounted for in the neuroimaging research, complementing the previous research regarding chronotype (the phase of the rhythms). We report that the structure of the early visual system is related to the perceived strength of mood and cognition fluctuations throughout the day independently of circadian preference. The presence of the opposite associations between the amplitude and the grey matter volume in the right middle temporal gyrus in the two chronotype groups indicates that the reported behavioural and neuronal chronotype differences might become more pronounced in individuals with extreme values of diurnal differences in mood and cognitive performance.

## Data availability

Data analysed in the current project is available online (Zareba et al., 2022).

## CRediT author statement

MRZ: Conceptualization, Methodology, Formal analysis, Writing - Original Draft, Visualization, Supervision. PS: Conceptualization, Methodology, Formal analysis, Writing - Original Draft. MF: Conceptualization, Investigation, Resources, Funding acquisition, Writing - review & editing. TM: Conceptualization, Funding acquisition, Writing - review & editing. HO: Conceptualization, Methodology, Resources, Writing - review & editing. IS: Writing - review & editing. EB: Conceptualization, Writing - review & editing. AD: Conceptualization, Investigation, Resources, Writing - review & editing.

## Ethical statement

The study was approved by the Ethics Committee of the Institute of Applied Psychology at Jagiellonian University.

